# The effect of spontaneous mutations on *CUP1* copy number and copper tolerance in *Saccharomyces cerevisiae*

**DOI:** 10.1101/2025.11.06.687044

**Authors:** Rohan Sohoni, Gillian G. Schatz, Nathaniel P. Sharp

## Abstract

Genomic regions containing tandem duplications may be subject to particularly high rates of copy number mutations due to recombinational repair of DNA damage. Consequently, changes in copy number may be among the most accessible beneficial mutations during evolution in a new environment. In the budding yeast *Saccharomyces cerevisiae*, the gene *CUP1* occurs in multiple tandem copies and encodes a metallothionein protein that is relevant for copper homeostasis. We examined *CUP1* copy number in 220 mutation accumulation lines and their ancestors to quantify spontaneous genetic change at this locus. We also measured copper tolerance in the same strains to understand the phenotypic effects of *CUP1* copy number change and other mutations. We found that mutations in *CUP1* copy number occurred rapidly, with a bias towards the loss of copies. While copper tolerance also declined due to spontaneous mutations, this change was less rapid overall and was only partly driven by *CUP1* copy number. Thus, while this locus is highly susceptible to mutation, such mutations may provide an accessible but insufficient evolutionary path towards increased copper resistance.

## Introduction

Different kinds of spontaneous mutations, ranging from single-nucleotide changes to whole-genome duplications, can occur at different rates and vary in their phenotypic impact. Gene duplications likely play an important role in genome evolution, but it is challenging to obtain measures of spontaneous duplication rates that are not influenced by selection [1–3]. There is evidence that tandem duplications are particularly susceptible to copy number mutation due to unequal mitotic recombination and other DNA double-strand break repair pathways [4–7], but the spontaneous rate and direction of such changes has not been well characterized. One way to obtain unbiased estimates of mutation rates is by conducting a mutation accumulation (MA) experiment, where multiple lineages are repeatedly bottlenecked to limit the influence of selection and mutations are identified by genome sequencing [8,9]. In these experiments, copy number changes can be detected bioinformatically, including by examination of relative sequencing coverage at a locus of interest [6,10–15]. Mutation accumulation lines also provide opportunities to study whether the genetic changes identified are associated with changes in phenotype.

In the budding yeast *Saccharomyces cerevisiae*, one locus that exhibits natural copy number variation is *CUP1* [6,16–18]. *CUP1* is a small gene encoding a metallothionein protein, which mediates copper ion homeostasis [19]. The laboratory strain S288c is reported to have 15 tandem copies of this gene, and a sample of natural isolates were found to have 2–18 copies [6]. The complete absence of *CUP1* is not lethal, but strains with one copy or less are described as copper sensitive [20]. There is evidence that strains with more copies of *CUP1* are more copper tolerant and that increased copy number often evolves in high-copper environments [6,17,20–23]. However, it is unclear how rapidly copy number changes spontaneously, and what the effects of un-selected copy number variation are on copper tolerance.

We examined copy number at the *CUP1* locus in 220 haploid and diploid lines of *S. cerevisiae* that each experienced about 1500 generations of MA in standard media [24], and performed growth inhibition assays to estimate relative copper tolerance in these same lines. Our main aims were to quantify the rate of mutational change in both *CUP1* copy number and copper tolerance, as well as the relationship between these traits. We also looked for point mutations within *CUP1* and tested whether other mutations known to be present in these strains [24] were associated with copper tolerance.

## Materials and methods

### Yeast strains

We conducted our study using the mutation accumulation (MA) lines and corresponding genome sequence data generated from Sharp et al. [24]. That experiment compared mutation in haploid and diploid versions of the strain SEY6211, where some lines of each ploidy type were genetically modified to delete the gene *RDH54*, which is involved in DNA double strand break repair. In our study, we considered possible effects of ploidy and *RDH54* status, but our main interest was in the consequences of mutation accumulation. We therefore compared MA lines with their ancestors of the same ploidy and *RDH54* status, which remained frozen at –80 C during the MA experiment.

### CUP1 copy number and point mutations

Our estimates of *CUP1* copy number are based on Illumina NextSeq data (150-bp paired-end reads), aligned to the reference genome as described in Sharp et al. [24]. Coverage-based estimates of *CUP1* copy number have been found to be strongly correlated with estimates by Southern blotting [6]. In the reference genome, the *CUP1* gene is represented by two copies at positions 212535-212720 and 214533-214718 on chromosome VIII. For each MA and ancestral strain, we used Samtools [25] to determine the average sequencing coverage at the *CUP1* sites, as well as the average coverage on chromosome VIII (using medians produced similar results). We divided *CUP1* coverage by chromosome VIII coverage and multiplied by two to obtain an estimate for *CUP1* copy number per chromosome VIII. This will reflect the copy number per cell in haploids, and half the copy number per cell in diploids, averaged over homologous chromosomes. In two diploid MA lines, chromosome VIII is known to be present in three copies (trisomy; [24]), and so we further multiplied our copy number estimates by 1.5 for these two lines when examining the relationship between copy number and copper tolerance.

In addition to copy number, mutations within the CUP1 gene could affect copper tolerance, but this region was excluded from a previous analysis of point mutations [24]. To detect point mutations in *CUP1*, we generated pileup files using Samtools [25] and then applied VarScan [26], with the minimum variant allele frequency threshold set to zero, allowing all variants to be reported, even those supported by a single read.

### Copper tolerance

We estimated copper tolerance based on growth inhibition on plates, blocking the MA lines by mating type (haploid MATa, haploid MAT*α*, or diploid) and *RDH54* status (wild type, *RDH54+* or deleted, *rdh54Δ*), with the corresponding ancestral control strains included in each block. For each block, we revived frozen strains in 96 well plates containing 500-750 µL of yeast peptone dextrose (YPD) medium per well, with 48 h of growth at 30 C. We tested copper tolerance on 6 cm diameter plates containing 8 mL of YPD-agar medium by spreading 50-75 µL of yeast culture over each plate using sterile beads. We then placed a sterile filter paper circle (diameter 2.3 cm) in the center of each plate and added 200 µL of copper sulfate pentahydrate in deionized water (312.5 g/L) to the filter circle. For each of the 220 MA lines we obtained three replicate measures of copper tolerance for a total of 657 replicates (growth inhibition zones were unclear on three plates, which were excluded). We also obtained 18 replicate measures of copper tolerance for each of the six types of ancestral control, for a total of 108 replicates. After 24 h of incubation at 30 C, we imaged the plates using a flatbed scanner. We used ImageJ [27] to determine the area of the zone of yeast growth inhibition around the filter circle on each plate (arbitrary units) and used the reciprocal of this value as an estimate of copper tolerance, i.e., a larger zone of growth inhibition equates to lower tolerance. We analyzed copper tolerance data using linear mixed-effect models fit in *R* [28] using packages lme4 [29] and RLRsim [30].

## Results

Our analysis of sequencing coverage in the ancestor strains of the MA experiment revealed an average *CUP1* copy number of 10.8 per haploid genome, somewhat less than the estimate of 15 for the reference strain S288c, but within the range observed in natural isolates [6]. We found no evidence for variation in copy number among the ancestral control samples sequenced (which vary in ploidy and *RDH54* status but should otherwise be identical; all *F* < 2.4, *P* > 0.17), and so we treated the mean copy number of those strains as representative of the ancestral condition of all of the MA lines. Mean *CUP1* copy number declined significantly during MA going from 10.8 in the ancestor to an average of 8.6 in the MA lines (linear model, effect of MA: *t* = –3.34, *P* = 9.81 × 10^−4^) and ranging from 3.4 to 18.5 (Fig. 1). We found no evidence that the rate of copy number decline differed between groups (model interaction terms: *P* > 0.66). The rates of change in mean *CUP1* copy number per generation (Δ*M*) are given in Table 1. On average, we find a net loss of one *CUP1* copy per 709 generations. For individual lines, the fastest observed changes in each direction were one loss per 201 generations and one gain per 194 generations. Since we have only one estimate of *CUP1* copy number for each MA line, we cannot formally determine the mutational variance for this trait, but we used the variance among control samples to approximate the error variance. The variance values for each group, following subtraction of the error variance and division by the number of cell divisions (Δ*V*), are given in Table 1.

**Table 1.** Trait change per generation of mutation accumulation. Rates of change in mean (Δ*M*), variance (Δ*V*), and coefficient of variation (CV) per generation for copper tolerance and *CUP1* copy number.

| Cell type | *RDH54* | Copper tolerance |  |  | *CUP1* copy number |  |  |
| --- | --- | --- | --- | --- | --- | --- | --- |
| | | $\Delta M (10^{-6})$ | $\Delta V (10^{-6})$ | CV (%) | $\Delta M (10^{-3})$ | $\Delta V (10^{-3})$ | CV (%) |
| MATa | *RDH54+* | -48.83 | 1.26 | 0.12 | -1.35 | 2.58 | 0.58 |
| MATa | *rdh54Δ* | -50.70 | 2.28 | 0.16 | -1.91 | 3.57 | 0.75 |
| MATα | *RDH54+* | -16.70 | 1.44 | 0.12 | -1.03 | 1.74 | 0.49 |
| MATα | *rdh54Δ* | -24.89 | 0.28 | 0.05 | -1.34 | 3.25 | 0.65 |
| Diploid | *RDH54+* | -39.78 | 1.96 | 0.15 | -1.54 | 1.12 | 0.40 |
| Diploid | *rdh54Δ* | -8.59 | 1.58 | 0.13 | -1.46 | 1.78 | 0.52 |

**Figure 1.**
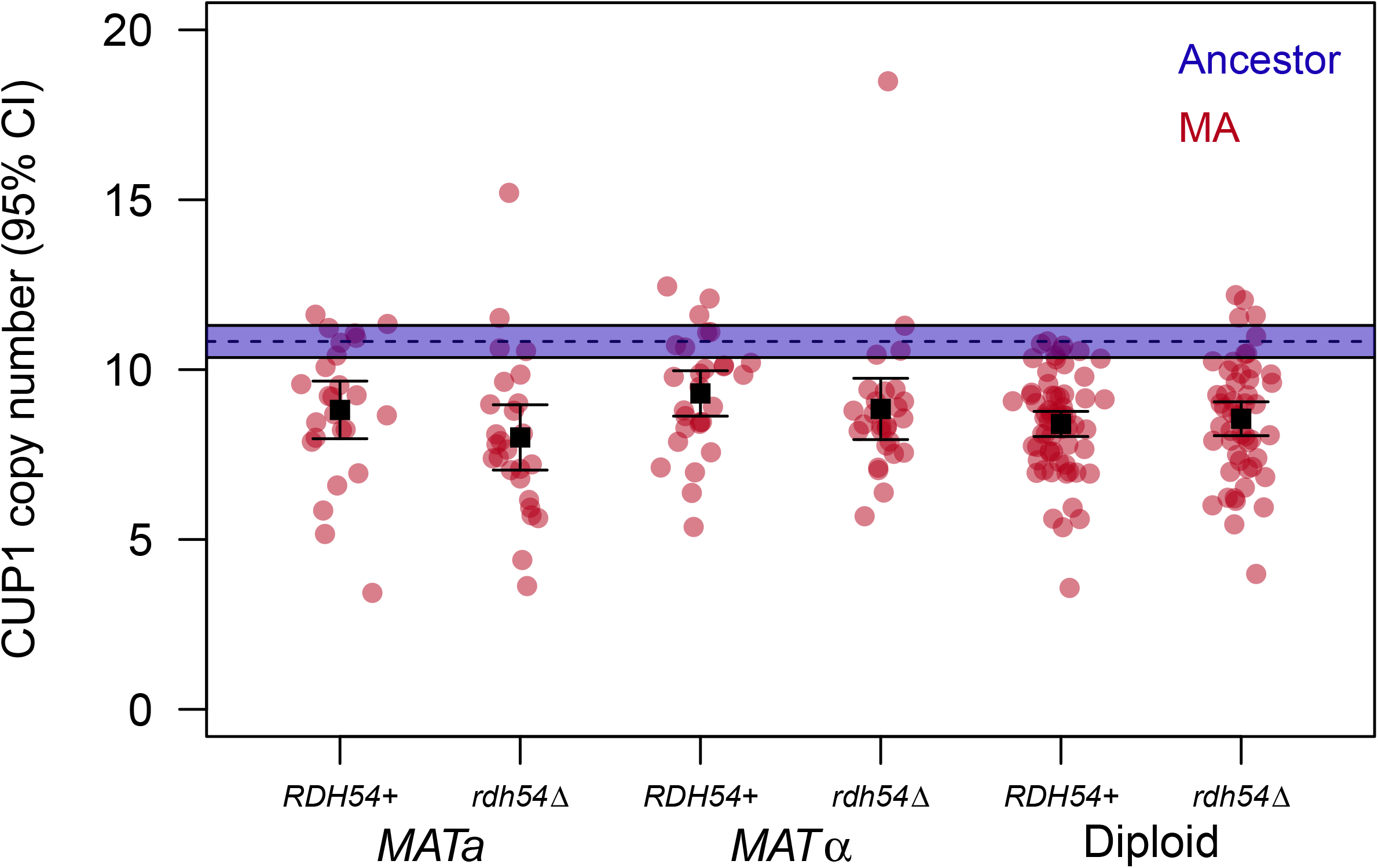
Mean *CUP1* copy number declines under MA. Each red point represents a *CUP1* copy number estimate for an MA line from a given cell type and *RDH54* status. Black squares and error bars represent group means and 95% confidence intervals. The mean and 95% confidence interval for the ancestral *CUP1* copy number is shown as a horizontal dashed line and band.

We found that average copper tolerance declined by about 5% during MA (about 0.003% per generation), with no evidence that this effect was related to cell type or *RDH54* status (Fig. 2; linear mixed-effect model, effect of MA: *t* = –4.04, *P* = 7.75 × 10^−5^; interactions with MA: *P* > 0.42). The rate of decline in copper tolerance per generation of MA (Δ*M*) in each group of lines is given in Table 1. We also tested for genetic variance in copper tolerance among MA lines within each group (mutational variance). We found strong evidence for mutational variance in all groups (all *P* < 10^−15^ except in MAT⌇0 *rdh54Δ* lines where *P* = 0.006). Rates of increase in mutational variance per generation of MA (Δ*V*) in each group of lines are given in Table 1.

**Figure 2.**
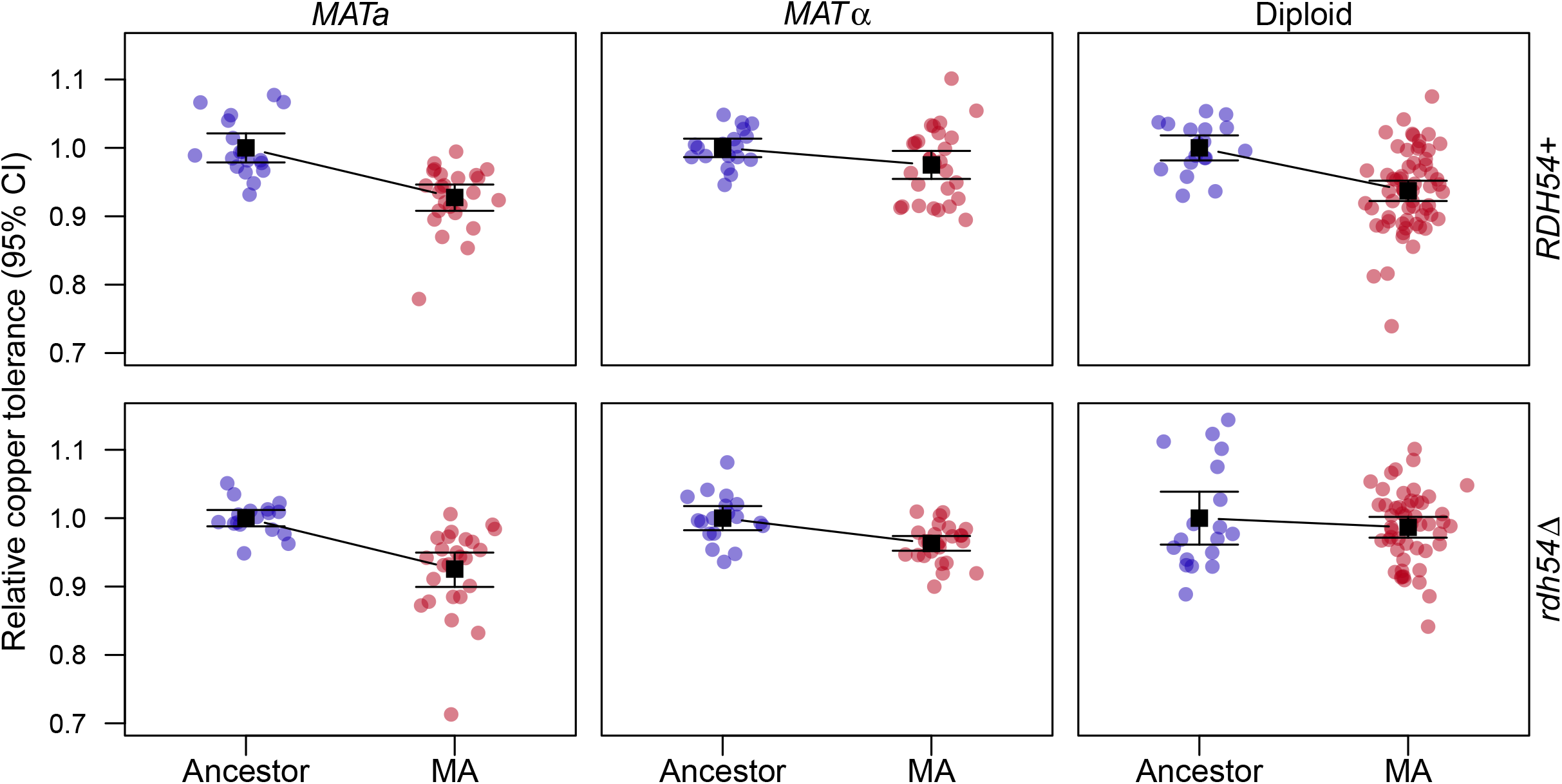
Mean copper tolerance declines under MA. Each round point represents a copper tolerance estimate for a given MA line or ancestral control replicate. Black squares and error bars represent group means and 95% confidence intervals. Panels depict groups of lines based on cell type and *RDH54* status.

We examined the relationship between copper tolerance and *CUP1* copy number, with tolerance in each MA line standardized by the corresponding ancestral control mean. We found a significant positive correlation between copper tolerance and *CUP1* copy number (Fig. 3A; *r*_Spearman_ = 0.21, *P* = 1.19 × 10^−3^), but with much variation in copper tolerance remaining unexplained by *CUP1* copy number (Pearson *r*^2^ = 2.6%). We also considered whether the rate of copy number change at *CUP1* might be related to the rate of copy number change at the highly repetitive rDNA locus, where copy number also declined under MA [14]. We found a significant positive correlation between *CUP1* and rDNA copy number (Fig. 3B; *r*_Spearman_ = 0.15, *P* = 0.028), suggesting that general mechanisms for copy number change may affect both loci. Decline in rDNA copy number was associated with point mutations in “regulation of cell cycle” genes [14], but we did not detect such an association for *CUP1* copy number (*r*_Spearman_ = 0.10, *P* = 0.15). A previous study of experimental evolution of copper tolerance found no evidence that increased *CUP1* copy number had pleiotropic costs for fitness in YPD [22]; we also found that *CUP1* copy number was not significantly correlated with growth rate in YPD (data from [24]; Fig. 3C; *r*_Spearman_ = –0.05, *P* = 0.44). Growth in YPD was also not significantly correlated with our measures of copper tolerance (*r*_Spearman_ = 0.01, *P* = 0.89).

**Figure 3.**
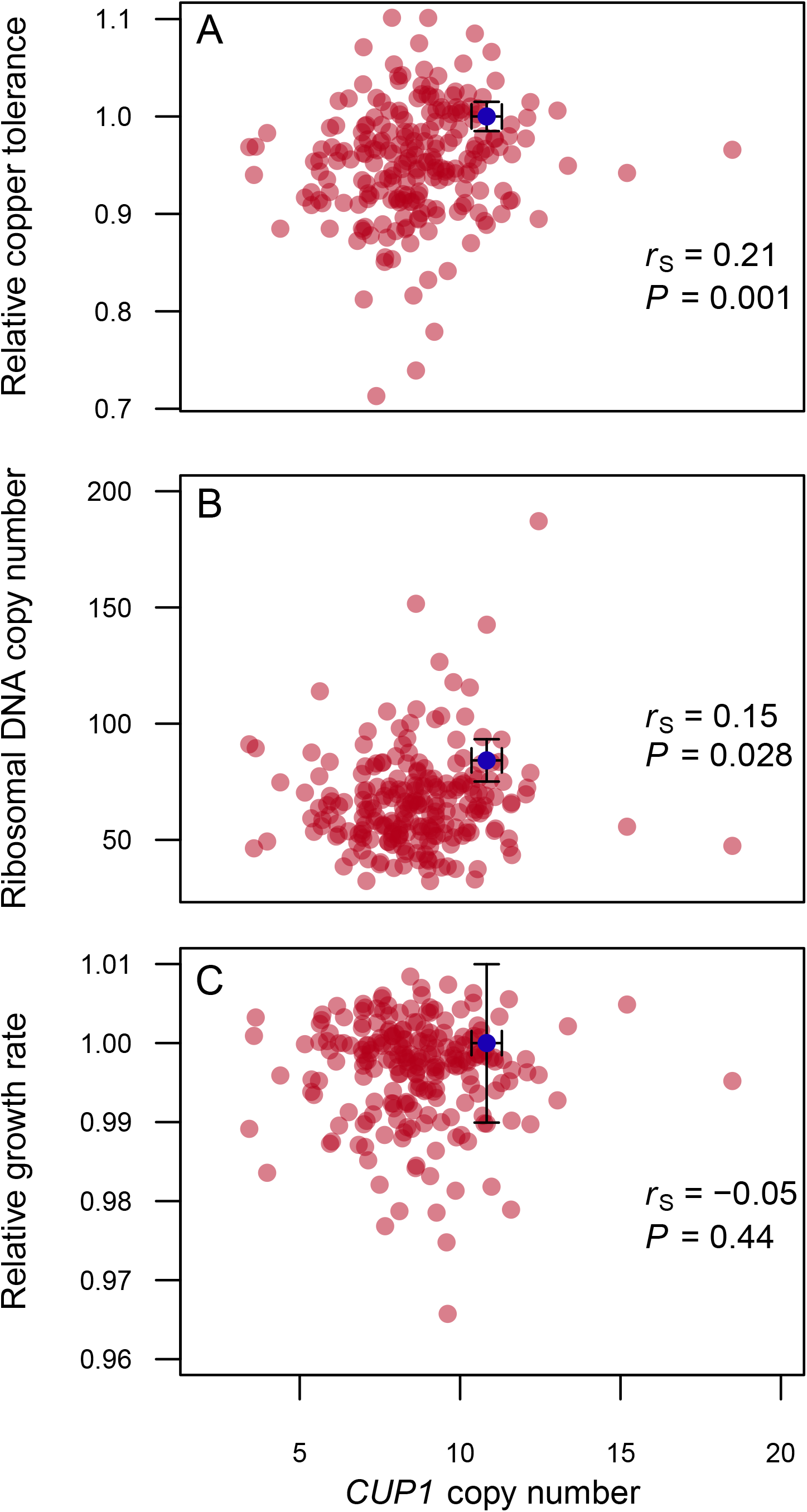
Associations between *CUP1* copy number and other traits. Each red point depicts mean trait values for a given MA line. The blue points and error bars represent the mean and 95% confidence interval for ancestral controls. The Spearman’s rank correlation (*r*_S_) and P-value (*P*) are shown in each panel. (A) *CUP1* copy number was positively associated with copper tolerance. (B) *CUP1* copy number was positively associated with rDNA copy number. (C) *CUP1* copy number was not significantly associated with growth rate relative to the ancestral control of the same cell type.

We did not detect any new point mutations or differences from the reference genome within the *CUP1* coding sequence. Given the point mutation rates in these strains [24], the number of MA generations, the size of *CUP1* (185 bp per copy), and assuming 8.6 copies per haploid genome (the post-MA mean), there is a 55% probability that no point mutations would occur in any *CUP1* copy during MA. Other genes are known to be upregulated in the presence of high copper, and additional genes have been found to acquire mutations under selection for copper tolerance [22,31]. The MA lines in our study accumulated un-selected mutations, which might affect these previously identified genes by chance. Two MA lines had distinct missense mutations in *PCA1* (A789T and A830P), which encodes a copper-binding protein that may be involved in metal ion homeostasis [32] but was not hit during experimental evolution [22]. These two strains had significantly below-average copper tolerance compared with the other MA lines (*t* = –8.27, *P* = 0.014), despite not differing from the average in terms of *CUP1* copy number (*t* = 1.14, *P* = 0.45). An additional five MA lines had nonsynonymous mutations in other genes implicated in copper tolerance evolution [22], but these lines did not show significantly reduced tolerance (*t* = –1.00, *P* = 0.36). We also noted that the MA line with the lowest observed copper tolerance (Fig. 3A) had a missense mutation in *FAB1* (T480I), which encodes a protein involved in vacuolar homeostasis, whose deletion confers metal ion sensitivity [33].

Increased copy number of chromosome II has also been implicated in copper tolerance, possibly because it carries several relevant genes [22,34]; here, six MA lines had extra copies of that chromosome, and we found suggestive evidence that these strains may have had higher than average copper tolerance (Wilcox test, *P* = 0.091). Considering aneuploidy more generally (36 of the MA lines are aneuploid for at least one chromosome), we found that chromosome gains were associated with greater copper tolerance (*r*_Spearman_ = 0.14, *P* = 0.033), whereas chromosome losses were not (*r*_Spearman_ = –0.01, *P* = 0.83). The association between tolerance and extra copies of each chromosome is shown in Table 2.

**Table 2.** Relative effects of chromosome gains on mean copper tolerance. Chromosomes where no aneuploidy was observed are omitted.

| Chromosome | N lines | Tolerance effect |
| --- | --- | --- |
| I | 7 | +1.2% |
| II | 6 | +3.94% |
| III | 4 | +0.71% |
| VI | 1 | +1.21% |
| VII | 2 | +9.51% |
| VIII | 2 | +5.19% |
| IX | 3 | +1.31% |
| X | 2 | +4.58% |
| XI | 2 | -2.05% |
| XII | 5 | -2.84% |
| XIII | 1 | -4.88% |
| XIV | 2 | +4.81% |

## Discussion

Although increased *CUP1* copy number seems to be common under experimental evolution [22], we find that there is a mutational bias towards the net loss of copies (Fig. 1). Of the 220 MA lines we examined, only 20 (9.1%) showed *CUP1* copy number greater than that of the ancestor, with just 6 lines (2.7%) showing increases of more than 1 copy. Additional copies of *CUP1* are likely to be beneficial in the presence of high levels of copper (Fig 3A; [6,20–23]), and there is no evidence that the presence of additional *CUP1* repeats has detrimental pleiotropic consequences for growth rate in the absence of copper (Fig. 3C; [22]). We would therefore expect selection to drive increased copy number in a high-copper environment, but for mutation bias and genetic drift to lead to reduced copy number in a low-copper environment.

The net rate of change in *CUP1* copy number per generation (Table 1) exceeds the genome-wide average rate of gene duplication or deletion in yeast and other model organisms by three orders of magnitude [2,35]. We are likely underestimating the true rate of mutational change because gains and losses in the same MA lineage will cancel out. In the genetic background we studied, *CUP1* is likely to be more susceptible to copy number change than most other genes because it is already present in multiple tandem copies, facilitating unequal crossing over [6]. In diploids, these events may be more likely to occur between sister chromatids rather than homologous chromosomes [36]; this is consistent with our observation that the deletion of *RDH54*, which is believed to facilitate DNA double-strand break repair between homologs [37] had no detectable effect on the rate of copy number mutation (Fig. 1).

The mutational properties of *CUP1* may be similar to those of the highly repetitive rDNA locus, which is present in a tandem array of about 90 copies in the genetic background we studied, and also seems to experience net copy number loss on average, with rarer cases of copy number gain [14]. Mutational change in rDNA is faster on an absolute basis, but when scaled to the initial copy number, both loci show an average decrease of about 0.01% per generation under mutation accumulation. We also detected a positive correlation across MA lines between *CUP1* copy number and rDNA copy number (Fig. 3B), suggesting that mutations influencing rDNA copy number maintenance affect other repetitive loci as well. Changes in rDNA were associated with point mutations in “regulation of cell cycle” genes [14], but we did not detect such an association for *CUP1* copy number.

In the strains we studied, copper tolerance showed a modest level of mutational variance [8,38]. While CUP1 copy number was a significant predictor of tolerance (Fig. 3A), there was plenty of unexplained variation. A trivial reason for this is that measurement error for both copper tolerance and *CUP1* copy number will reduce the apparent correlation between these traits (attenuation), but correcting for this bias would have only a small effect on the correlation in this case. Karyotypic changes and mutations at other loci may also have contributed to variation in tolerance (on average, each line had 5.2 non-synonymous mutations). There could also be diminishing returns of additional *CUP1* copies if transcripts from high-copy loci interfere with one another [39]. In natural populations, any initial change in transcript abundance following a gene duplication mutation could be modified by subsequent selection on other loci [2,12]. A previous study found weak evidence that *CUP1* expression was related to copy number in experimentally evolved strains [22].

Our study demonstrates that *CUP1* copy number can mutate rapidly, with a bias towards the loss of copies, as expected given its genome architecture. Despite rapid change at *CUP1*, the rate of mutational decline in copper tolerance was modest. While changes in *CUP1* copy number affected copper tolerance, point mutations in other genes were also associated with reduced tolerance, and copy number increases for several chromosomes were associated with increased tolerance. Increased *CUP1* copy number is likely to be an early but insufficient response to selection for copper tolerance.

## Data availability

Raw data and analysis code are available at https://doi.org/10.6084/m9.figshare.30559424.v1.

